# DNA methylation signature of smoking in lung cancer is enriched for exposure signatures in newborn and adult blood

**DOI:** 10.1101/356485

**Authors:** K. M. Bakulski, J. Dou, N. Lin, S. J. London, J. A. Colacino

## Abstract

**Background:** Smoking impacts DNA methylation genome-wide in blood of both newborns from maternal smoking during pregnancy and adults from personal smoking. Smoking causes lung cancer which involves aberrant methylation. We examined whether DNA methylation smoking signatures identified in blood of newborns and adults are detectable in lung tumors.

**Methods:** We compared smoking-related DNA methylation in lung adenocarcinomas (61 never smokers, 91 current smokers, and 238 former smokers) quantified with the Illumina450k BeadArray in The Cancer Genome Atlas with published large consortium meta-analyses of newborn and adult blood. We assessed whether CpG sites related to smoking in blood from newborns and adults were enriched in lung adenocarcinoma.

**Results:** Testing CpGs differentially methylated by smoke exposure (*P*<10^−4^) we identified 296 in lung tumors, while previous meta-analyses (False Discovery Rate (FDR)<0.05) identified 6,073 in newborn blood, and for adult smoking, 18,760 in blood. The lung signals were highly enriched for those seen in newborn (32 overlapping, P_*enrichment*_=1.2×10^−19^) and adult blood (86 overlapping, *P*_*enrichment*_ = 9.5×10^−49^). The 65 genes annotated to CpGs differentially methylated in lung tumors, but not blood, were enriched for RNA processing ontologies.

**Conclusions:** We found highly significant overlap between smoking-related DNA methylation signals in lung cancer and those seen in blood from newborns, from *in utero* exposure, or adults, from their own exposure. These results suggest that some epigenetic alterations associated with cigarette smoke exposure are tissue specific, but others are common across tissues. These findings support the value of blood-based methylation biomarkers for assessing exposure effects in target tissues.

## INTRODUCTION

Approximately one quarter of cancer deaths are attributable to tobacco use (1). The lung is the primary tissue affected by tobacco smoke and tobacco accounts for 87% of deaths due to lung cancer (1). Specifically, lung adenocarcinoma is the leading cause of cancer deaths globally. Epigenetic modifications, including DNA methylation, are widely detected in cancers including lung adenocarcinoma and may play a role in pathogenesis (2).

Exposure to cigarette smoke is associated with altered DNA methylation at many locations throughout the genome. A recent epigenome-wide meta-analysis of blood DNA methylation using the Illumina450K Beadchip in 6,685 newborns from 13 studies in the Pregnancy and Child Epigenetics (PACE) consortium identified over 6,000 CpG sites differentially methylated in relation to maternal smoking during pregnancy (3). Differential blood DNA methylation in relation to maternal smoking during pregnancy was subsequently shown to be a reliable biomarker of this exposure in newborns (4). In adults, personal smoking was related to widespread differential methylation in blood in a meta-analysis of 16 cohorts (n=15,907) in the Cohorts for Heart and Aging Research in Genetic Epidemiology (CHARGE) consortium (5). Exposure to cigarette smoke is associated with reproducible and specific DNA methylation changes in newborn and adult blood.

While blood is readily available in large scale population studies, the target tissues for the diseases of interest are not. A few studies have compared blood smoking candidate gene DNA methylation associations to lung, the primary organ exposed to smoke and the major target of smoking related carcinogenesis. In a subset of the European Prospective Investigation into Cancer and Nutrition (n=374), two candidate CpGs in the Aryl-Hydrocarbon Receptor Repressor (*AHRR*) gene were differentially methylated in relation to smoking in blood as well as differentially methylated in human lung tissue, with the same direction of effect. (6). An epigenome-wide association study of tobacco smoke exposure in lung tissue found eight CpG sites with reduced methylation in smokers, five of which had been previously identified in studies examining blood DNA methylation in relation to smoking (7). However, the smoking signatures from newborns or adults have not been systematically compared with signatures in a highly relevant target tissue for smoking related health effects, the lung.

Using published meta-analysis results from the PACE and CHARGE consortia, we compared smoking-related DNA methylation signatures detectable in blood to those in a well-characterized collection of lung adenocarcinoma cases from the Cancer Genome Atlas (TCGA). We sought to test whether the blood-based DNA methylation smoking signals at birth, from *in utero* exposure, and in adulthood, from personal exposure, are reflective of smoking-related methylation in lung tumor tissue.

## METHODS

### Lung adenocarcinoma study sample

The current study analyzed publicly available data from The Cancer Genome Atlas (TCGA). A total of 507 lung adenocarcinoma samples were obtained at surgery from individuals with lung adenocarcinoma (2). Smoking status was assessed by questionnaire (never, current, former >15 years, former ≤15 years). DNA was extracted and bisulfite converted as previously described (2). DNA methylation was measured using the Illumina Infinium HumanMethylation450 BeadChip Kit (450k) (8), a validated tool for quantifying genome-scale DNA methylation (9). Lung adenocarcinoma samples were interspersed across 20 plates with samples from other tissues.

### Lung DNA methylation data preprocessing

Raw methylation image files were downloaded from the Genomic Data Commons (GDC). We calculated and analyzed methylated (M) and unmethylated (U) intensities for low-quality samples (M<11, U<11) (n=37). Samples were removed if more than 1% of probes did not meet a detection p value of 0.01 (n=43). Probes with low detection (*P* < 0.01 in >10% of samples, n=4,043) and cross-reactive probes (n=29,233) were removed (10). Some individuals had multiple samples with methylation measures (n=36). For these individuals, we selected the sample with the smallest proportion of probes failing detection p-value. Normal-exponential using out-of-band probe (noob) background correction was used (11).

### Relation of lung DNA methylation to smoking

We tested for an association between DNA methylation and categorical smoking status, either current or former (recent ≤15 year quitters and longer >15 year quitters as separate groups) with reference to never smoking using multivariable linear regression in limma (12), with empirical Bayesian standard error adjustment (13). We adjusted for sex, age, cancer stage, plate, and the first ten principal components (PCs) of ancestry (14). In a sensitivity model, we used surrogate variable adjustment (sva) to adjust for unmeasured confounding and potential batch effects (15). We generated plots of observed versus expected *P* values and calculated lambda genomic inflation (16). We flagged “gap probes” that have clustered methylation distributions, likely due to underlying genetic variation, using the gap hunter function in minfi with 5% as the distance threshold to define gaps and 1% as the outlier group cutoff (17).

### Smoking signatures in newborn and adult blood

Published results from previous, large epigenome-wide association meta-analyses were used to define smoking related blood DNA methylation signatures. In the Pregnancy And Childhood Epigenetics (PACE) consortium, maternal sustained smoking (not including women who quit early in pregnancy) was associated with newborn blood 450k DNA methylation in 13 cohorts (3). Meta-analysis results for the 6,073 CpGs significantly related to maternal smoking (False Discovery Rate (FDR) q value < 0.05, corresponding to *P*<6.5×10^−4^) were available. Of these 6,073, data were available for 5,936 in the analysis of lung adenocarcinoma. In the Cohorts for Heart and Aging Research in Genomic Epidemiology (CHARGE) consortium, current smoking status was associated with adult blood 450k DNA methylation in 16 cohorts (5). Results from the 18,760 probes significantly related to smoking (FDR q value < 0.05, corresponding to *P* < 1.9×10^−3^) in CHARGE were available. Of these 18,760smoking associated probes, data were also available for 18,126 probes in the lung adenocarcinoma analyses. We used site-specific smoking effect estimates, standard errors, and p-values from each of the two studies.

### Enrichment testing

Our primary analysis of DNA methylation in lung adenocarcinoma compared current smokers to never smokers. We first tested all pairwise Pearson correlations between effect estimates or p-values for the newborn blood sustained smoke exposure, adult current smoking, and lung current smoking results. We examined enrichment of the blood smoking signatures in the lung results by looking at the overlap of CpG sites that met significance thresholds. We varied the p-value cutoff for inclusion of probes in the lung smoking signature from 1.0 to 10^−10^ in order to evaluate sensitivity to our choice of threshold. The threshold for the newborn smoke exposure signature and the adult blood signature was FDR<0.05. Fisher’s exact tests were used to determine significance of the overlap between lung and blood associated CpGs. We used the Illumina 450k annotation file to compare enriched sites by genomic region (CpG island, shore, shelf, or open sea).

### Gene ontology analysis

We examined the annotated genes of top differentially methylated CpG sites that were uniquely differentially methylated in lung adenocarcinoma of current smokers for enriched gene ontologies. We further tested for enriched ontologies from genes annotated to smoking-associated CpG sites that overlapped between lung adenocarcinoma and either newborn or adult blood. A *P* cut-off of 10^−4^ in lung and FDR cut-off of 0.05 for blood were the criteria used to determine CpG sites for inclusion in gene ontology analysis using the missMethyl package (18). We compared ontologies findings from the two sets of analyses performed: one considering genes with differentially methylated sites in lung only, and second that considered genes with differentially methylated sites in both lung adenocarcinoma and blood. Restricting to biological pathways with more than five genes, REVIGO was used to remove redundant gene ontologies (19).

### Secondary analyses

We tested for association in lung adenocarcinoma among former smokers at two different durations of time since quitting smoking (≤15 years and >15 years) relative to never smokers. We repeated the correlation and enrichment testing as specified above. As a sensitivity test, we also repeated enrichment analysis using a *P*<10^−4^ cutoff in the blood.

### RESULTS

#### Lung adenocarcinoma study sample descriptive statistics

After preprocessing, DNA methylation data were available on lung adenocarcinomas from 423 individuals in the TCGA. We analyzed the 390 participants with data on self-reported smoking status and covariates of interest (**Table 1**). The study sample was 54.9% female, 82.3% White, and 55.4% had cancer stage 1. Participants were a mean age of 65.3 at surgery. Smoking status differed by sex and age (**Table 2**).

**Table 1.**
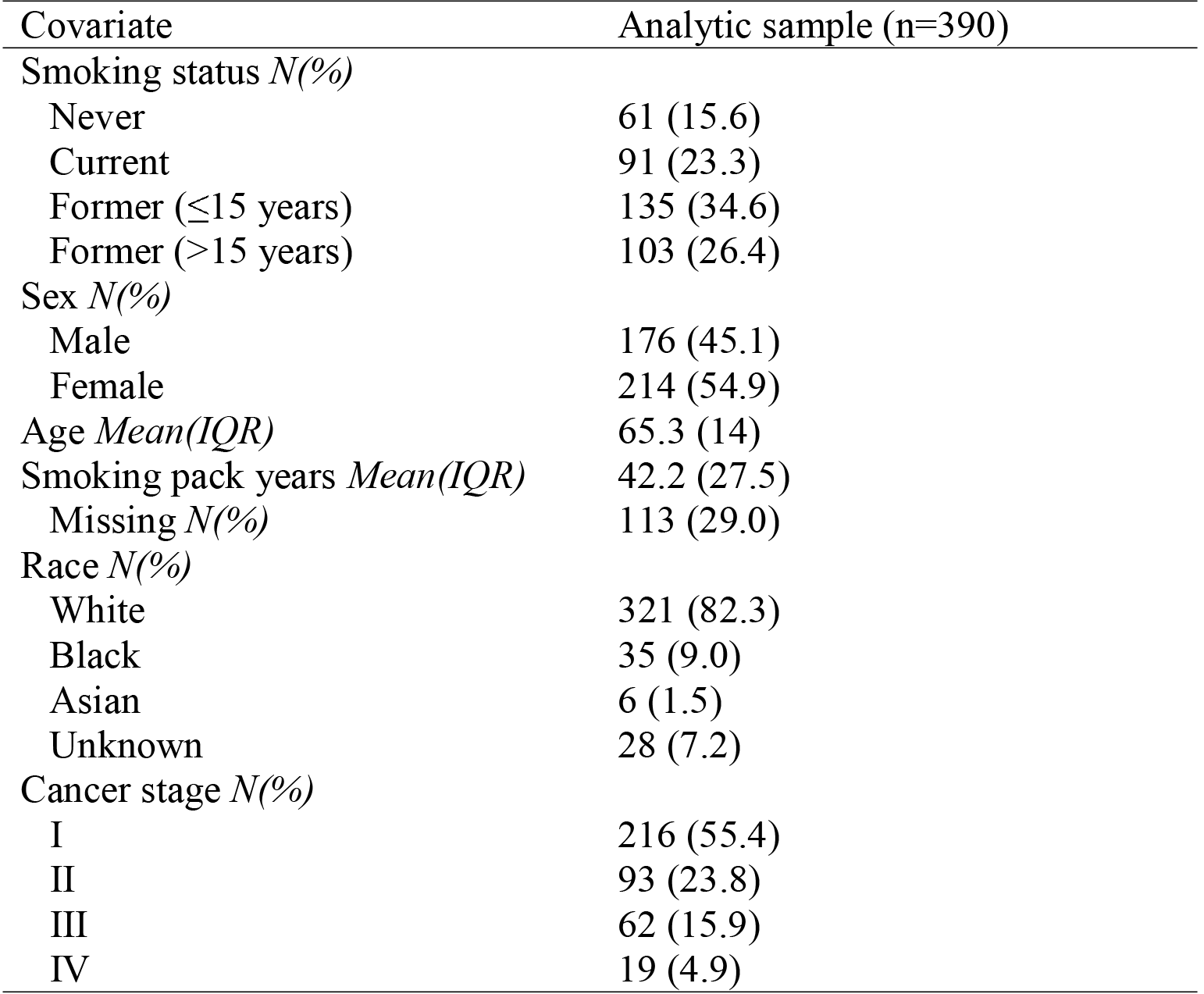
Univariate descriptive statistics of The Cancer Genome Atlas lung adenocarcinoma tissue study sample. We report mean(IQR) for continuous covariates and frequency (percent) for categorical covariates.

| Covariate | Analytic sample (n=390) |
| --- | --- |
| Smoking status *N*(%) |  |
| Never | 61 (15.6) |
| Current | 91 (23.3) |
| Former ($\leq 15$ years) | 135 (34.6) |
| Former ($> 15$ years) | 103 (26.4) |
| Sex *N*(%) |  |
| Male | 176 (45.1) |
| Female | 214 (54.9) |
| Age *Mean*(*IQR*) | 65.3 (14) |
| Smoking pack years *Mean*(*IQR*) | 42.2 (27.5) |
| Missing *N*(%) | 113 (29.0) |
| Race *N*(%) |  |
| White | 321 (82.3) |
| Black | 35 (9.0) |
| Asian | 6 (1.5) |
| Unknown | 28 (7.2) |
| Cancer stage *N*(%) |  |
| I | 216 (55.4) |
| II | 93 (23.8) |
| III | 62 (15.9) |
| IV | 19 (4.9) |

**Table 2.**
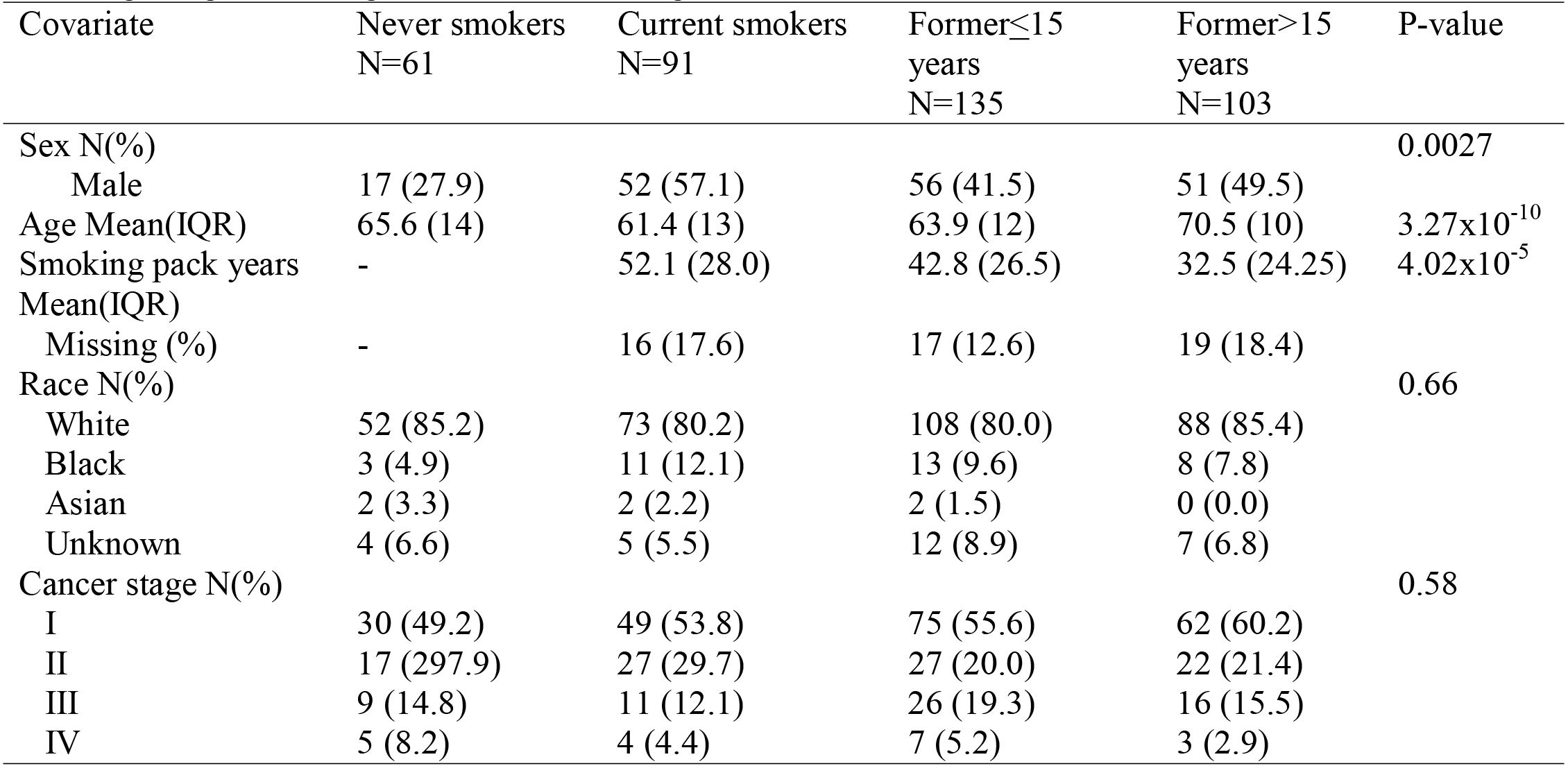
Bivariate descriptive statistics in the lung adenocarcinoma study (N=390). We report mean (IQR) for continuous covariates and frequency (percent) for categorical covariates and test for differences among the smoking categories, using Fisher’s test for categorical variables and ANOVA for continuous variables.

#### Smoking status and DNA methylation in lung adenocarcinoma samples

Our primary lung adenocarcinoma models included adjustment for age, 10 ancestry principal components (PCs), batch, cancer stage, and sex. For the primary model of current smoking versus never smoking in relation to DNA methylation we did not observe major inflation (lambda=1.13) (**Supplementary Figure 1**). The lambda value was slightly reduced by adding 10 surrogate variables to the primary model (current smoker lambda=1.08). In our primary model, comparing to never smokers, we observed lambda values of 1.05 for recent former smokers and 0.99 for longer-term former smokers (**Supplementary Figure 2**). Beta coefficients in the surrogate variable model and the primary model had Pearson correlation of 0.90 (**Supplementary Figure 3**). Results across all probes can be found in **Supplementary Table 1**.

In lung adenocarcinoma samples, there were 14 CpGs associated with current smoking status at a Bonferroni adjusted genome-wide significance level (*P*<10^−7^) (**Table 3**), 66 CpGs associated with current smoking stats at FDR significance (**Supplemental Table 2**), and 34,795 CpGs associated at a nominal level (*P*<0.05) (**Supplementary Figure 4**). The genome-wide significant CpGs annotated to multiple small nucleolar RNA genes, ribosomal subunit genes (*RSPs*), myosin immunoglobulin (*MYO1G*), and zinc finger protein 28 (*ZFP28*). Both CpGs annotated to *RPS8*, cg13985198 and cg18806997, were also FDR significant in adult blood (5) and cg13985198 was additionally significant in newborn blood (3). Both genome-wide significant CpGs in *RPS18*(also mapping to vacuolar protein sorting 52 (*VPS52*)), cg07362537 and cg12086028, were FDR significant in both adult and newborn blood (3, 5). Significant CpGs in *MYO1G,* cg22132788 and cg1908920, were also found in adult and newborn blood (3, 5).

**Table 3.** CpG sites with genome-wide significance level (*P* value <10^−7^) for either current or former smoking (versus never smoking) in The Cancer Genome Atlas lung adenocarcinoma samples.

| CpG | Chr | Position | Annotated Genes | Estimated Difference in % Methylation | Std. Error | P-value | Average % Meth. | Gap Probe |
| --- | --- | --- | --- | --- | --- | --- | --- | --- |
| **Current Smoking** |  |  |  |  |  |  |  |  |
| cg12086028 | chr6 | 33241026 | VPS52; RPS18 | -15.12 | 2.37 | 5.53E-10 | 31.5 | N |
| cg27033919 | chr11 | 62622173 | SNORD30; SNORD29;<br>SLC3A2; SLC3A2; SNORD31;<br>SNORD28; SLC3A2; SLC3A2;<br>SNHG1; SLC3A2 | -13.19 | 2.16 | 2.79E-09 | 13.4 | Y |
| cg18806997 | chr1 | 45242078 | SNORD46; RPS8; SNORD38A | -7.2 | 1.19 | 3.34E-09 | 9.6 | N |
| cg22132788 | chr7 | 45002486 | MYO1G | 13.31 | 2.24 | 7.16E-09 | 76.3 | N |
| cg16290996 | chr1 | 173835989 | GAS5; SNORD78; SNORD76;<br>SNORD77; SNORD44;<br>SNORD79 | -12.64 | 2.13 | 7.26E-09 | 21.8 | N |
| cg13985198 | chr1 | 45242073 | SNORD46; RPS8; SNORD38A | -8.22 | 1.39 | 7.63E-09 | 6 | Y |
| cg02607319 | chr7 | 45002112 |  | 12.95 | 2.23 | 1.36E-08 | 69.2 | N |
| cg07362537 | chr6 | 33240820 | VPS52; RPS18 | -13.18 | 2.35 | 3.96E-08 | 21.2 | Y |
| cg02905828 | chr11 | 62622234 | SNORD30; SNORD29;<br>SLC3A2; SLC3A2; SNORD31;<br>SNORD28; SLC3A2; SLC3A2;<br>SNHG1; SLC3A2 | -13.25 | 2.37 | 4.36E-08 | 16.5 | N |
| cg11374355 | chr19 | 39925660 | RPS16 | -9.26 | 1.66 | 5.27E-08 | 42.1 | N |
| cg12973930 | chr19 | 57049695 | ZFP28 | -10.66 | 1.92 | 5.73E-08 | 11.5 | N |
| cg19089201 | chr7 | 45002287 | MYO1G | 11.88 | 2.16 | 7.42E-08 | 68 | N |
| cg09345320 | chr11 | 62622179 | SNORD30; SNORD29;<br>SLC3A2; SLC3A2; SNORD31;<br>SNORD28; SLC3A2; SLC3A2;<br>SNHG1; SLC3A2 | -13.27 | 2.43 | 8.39E-08 | 13 | Y |
| cg05840553 | chr9 | 130212635 | RPL12; LRSAM1; LRSAM1 | -6.28 | 1.15 | 8.52E-08 | 6.8 | Y |
| **Former Smoking ($\leq 15$ Years)** |  |  |  |  |  |  |  |  |
| cg22132788 | chr7 | 45002486 | MYO1G | 15.03 | 2.05 | 1.57E-12 | 76.3 | N |
| cg02607319 | chr7 | 45002112 |  | 14.79 | 2.03 | 2.31E-12 | 69.2 | N |
| cg19089201 | chr7 | 45002287 | MYO1G | 13.17 | 1.97 | 9.73E-11 | 68 | N |
| cg04180046 | chr7 | 45002736 | MYO1G | 12.01 | 1.91 | 1.03E-09 | 57.8 | N |
| cg23160522 | chr15 | 75015787 | CYP1A1 | -8.68 | 1.44 | 4.34E-09 | 57.9 | N |
| cg18806997 | chr1 | 45242078 | SNORD46; RPS8; SNORD38A | -6.12 | 1.08 | 3.34E-08 | 9.6 | N |
| cg13985198 | chr1 | 45242073 | SNORD46; RPS8; SNORD38A | -7.14 | 1.27 | 3.69E-08 | 6 | Y |
| cg18092474 | chr15 | 75019302 | CYP1A1 | -12.32 | 2.21 | 4.65E-08 | 43.6 | N |
| cg01410359 | chr2 | 38302230 | CYP1B1 | -10.14 | 1.82 | 4.66E-08 | 33.5 | N |
| cg03224163 | chr7 | 139420300 | HIPK2; HIPK2 | -3.89 | 0.7 | 6.01E-08 | 83.1 | Y |
| cg09799983 | chr2 | 38301756 | CYP1B1 | -23.47 | 4.25 | 6.59E-08 | 39.3 | N |
| cg12803068 | chr7 | 45002919 | MYO1G | 9.37 | 1.72 | 9.84E-08 | 73.4 | N |
| **Former Smoking ($> 15$ Years)** |  |  |  |  |  |  |  |  |
| cg25157280 | chr3 | 48700498 | CELSR3 | -8.97 | 1.57 | 2.27E-08 | 5.9 | N |
| cg22132788 | chr7 | 45002486 | MYO1G | 11.88 | 2.17 | 8.43E-08 | 76.3 | N |
| cg09736162 | chr3 | 48700443 | CELSR3 | -9.35 | 1.71 | 9.34E-08 | 9.1 | N |

Second, we tested lung adenocarcinoma samples for differences between recent former smokers (≤15 years) and never smokers. There were 12 CpGs associated with recent former smoking status at a genome-wide significance level (*P*<10^−7^), with multiple CpGs in *MYO1G* and cytochrome p450 family genes *CYP1A1* and *CYP1B1*. Interestingly, all three of these genes were differentially methylated at genome wide significance in blood in relation to smoking in both newborns and adults (3, 5). The genome-wide significant CpG annotated to *HIPK2* (cg03224163) was also found in adult blood (5). There were 19 sites associated with recent former smoking at FDR<0.05 (**Supplemental Table 2**) and 29,480 CpGs associated at a nominal significance level (P<0.05).

Last, we tested for differences between longer term former smokers (> 15 years) and never smokers in the lung adenocarcinoma samples. In former smokers quitting >15 years ago, there were three genome-wide (*P*<10^−7^) significant sites, 14 FDR significant sites (**Supplementary Table 2**), and 24,620 CpGs were associated with longer former smoking at a nominal level (*P*<0.05). There were two CpGs associated with longer former smoking status at a genome-wide level annotated to cadherin EGF LAG Seven-Pass G-Type Receptor 3 (*CELSR3*), a gene identified previously in adult blood of former smokers relative to never smokers (5). The other significant CpG site, cg22132788, was mapped to *MYO1G* and was one of the same sites found in both recent former smokers and current smokers.

#### Correlation of CpGs differentially methylated in related to smoking in adult blood, newborn blood, and lung adenocarcinoma

We next sought to compare the overall pattern of smoking related DNA methylation sites pairwise across the three tissues: adult blood, newborn blood, and lung adenocarcinoma. First, there were 1,376 CpG sites that were FDR significant (FDR<0.05) in both the adult blood and the newborn blood meta-analyses. At these sites, adult blood and newborn blood smoking effect estimates were highly correlated (Pearson r=0.57, P<1.1×10^−16^, n=1,376). Across the 5,936 FDR significant CpG sites in newborn blood that were present in the adenocarcinoma dataset at any level of significance, the effect estimates describing the relationship between current versus never smoking in lung adenocarcinoma and estimates in newborn blood for maternal smoking in pregnancy were only weakly, albeit significantly, correlated (Pearson r=0.04, *P*=0.0034). For FDR significant (FDR<0.05) probes in adult blood, there was slightly stronger correlation between effect estimates for personal smoking in adult lung and blood (Pearson r=0.13, P<1.1×10^−16^, n=18,126) (**Supplementary Figure 5**). A similar pattern held for the–log10 *P* values where analogous correlations for differential methylation findings in lung and blood were slightly stronger in adult than in newborn blood (Pearson r=0.20 for lung and adult blood, r=0.08 for lung and newborn blood, both P<1.1×10^−16^).

#### Lung adenocarcinoma smoking DNA methylation signature is enriched for adult blood smoking DNA methylation signature

When we compared the individual CpG sites associated with smoking in lung adenocarcinoma and adult blood, 18,126 CpG sites associated with current smoking in adult blood that met FDR significance and were represented in the lung adenocarcinoma dataset. We tested the differentially methylated CpGs in lung for enrichment of those signals in adult blood using variable cutoffs in lung adenocarcinoma results (**Figure 1**), and here we report results using a *P*<10^−4^ threshold in lung. Among CpG sites distinguishing current smokers versus never smokers in lung adenocarcinoma at *P*< 10^−4^ (n=296 sites), we observed highly significant enrichment for the adult blood signature at *P* = 9.5×10^−49^. This corresponded to an observed overlap of 86 sites, relative to an expected overlap of 11.8 sites. Lung and adult blood current smoking effect estimates for the 86 overlapping sites had Pearson correlation of 0.36 (**Figure 2A**). A majority (75.6%) of these sites were hypomethylated in relation to smoking in both lung and adult blood. Among the sites with the highest absolute difference in methylation in both lung and adult blood were cg05575921 (*AHRR*) with decreased methylation in relation to smoking (lung: 9.6%, adult blood: 18.0%) and cg12803068 (*MYO1G*) with increased methylation (lung: 8.4% higher, adult blood: 6.3%). Among the sites overlapping between the lung adenocarcinoma and adult blood, the distribution of probes across genomic regions in relation to CpG islands differed from the full array distribution (G-test of goodness of fit *P*=6.2×10^−19^). We observed that 65.1% of overlapping sites were located in shore regions compared to 23.3% in the full array (*P*<2.2×10^−16^), 8.1% in open sea compared to 35.7% (*P*=5.5×10^−8^), and 19.8% in islands compared to 31.6% (*P*=0.026), while the proportion of overlapping sites in shelves (8.1%) was similar to the 9.4% in the full array (P=0.79).

**Figure 1.**
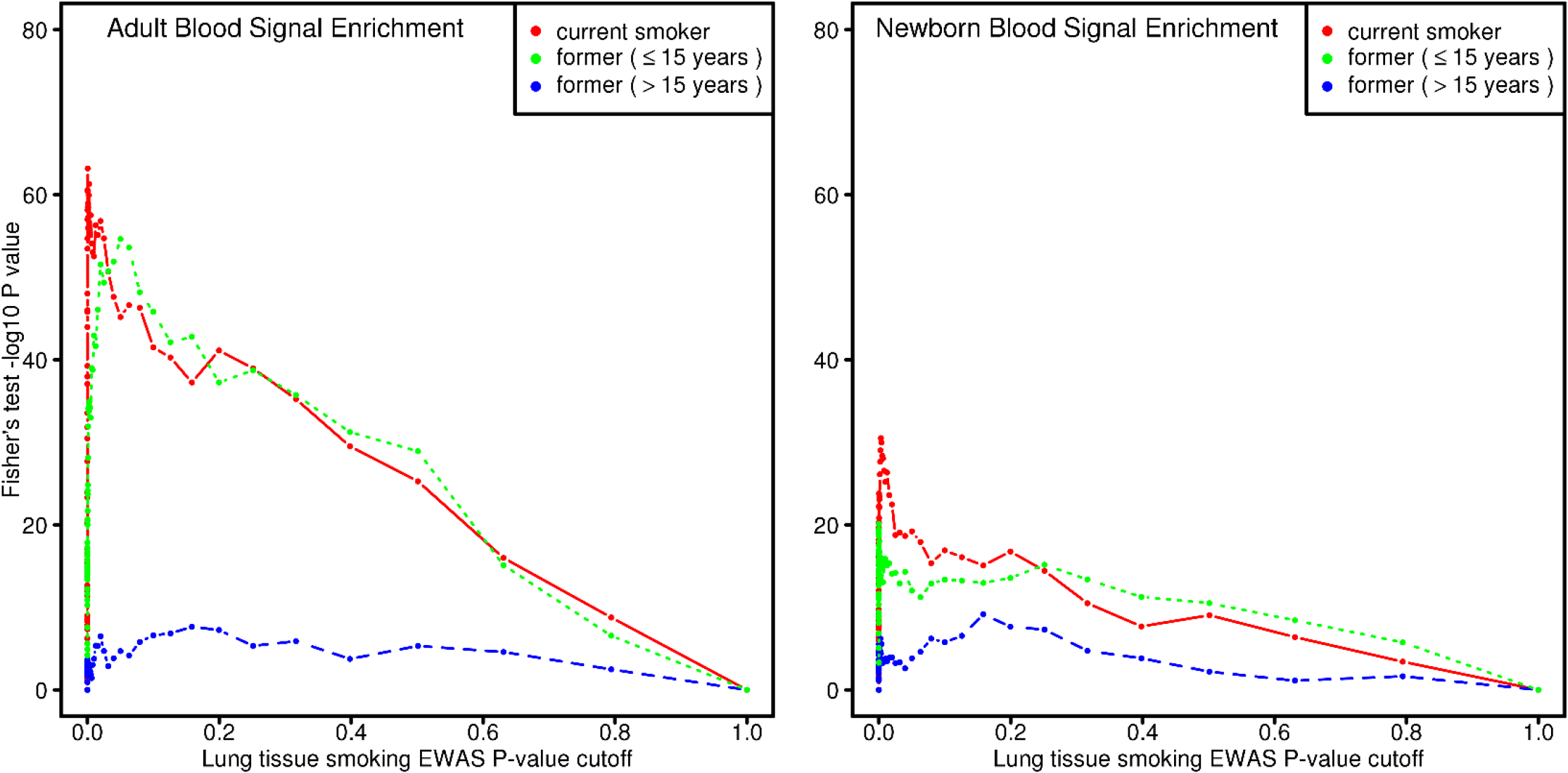
Enrichment for smoking associated DNA methylation sites in blood were evaluated by testing the overlap with smoking associated sites in lung adenocarcinoma of current smokers (red), recent former smokers (green), and longer former smokers (blue) that met P thresholds. The threshold in adult and newborn blood meta-analyses was an FDR<0.05. In lung, a range of P thresholds from 0 to 1 were applied. Fisher’s test was done for each P threshold on the overlap in CpG sites.

**Figure 2.**
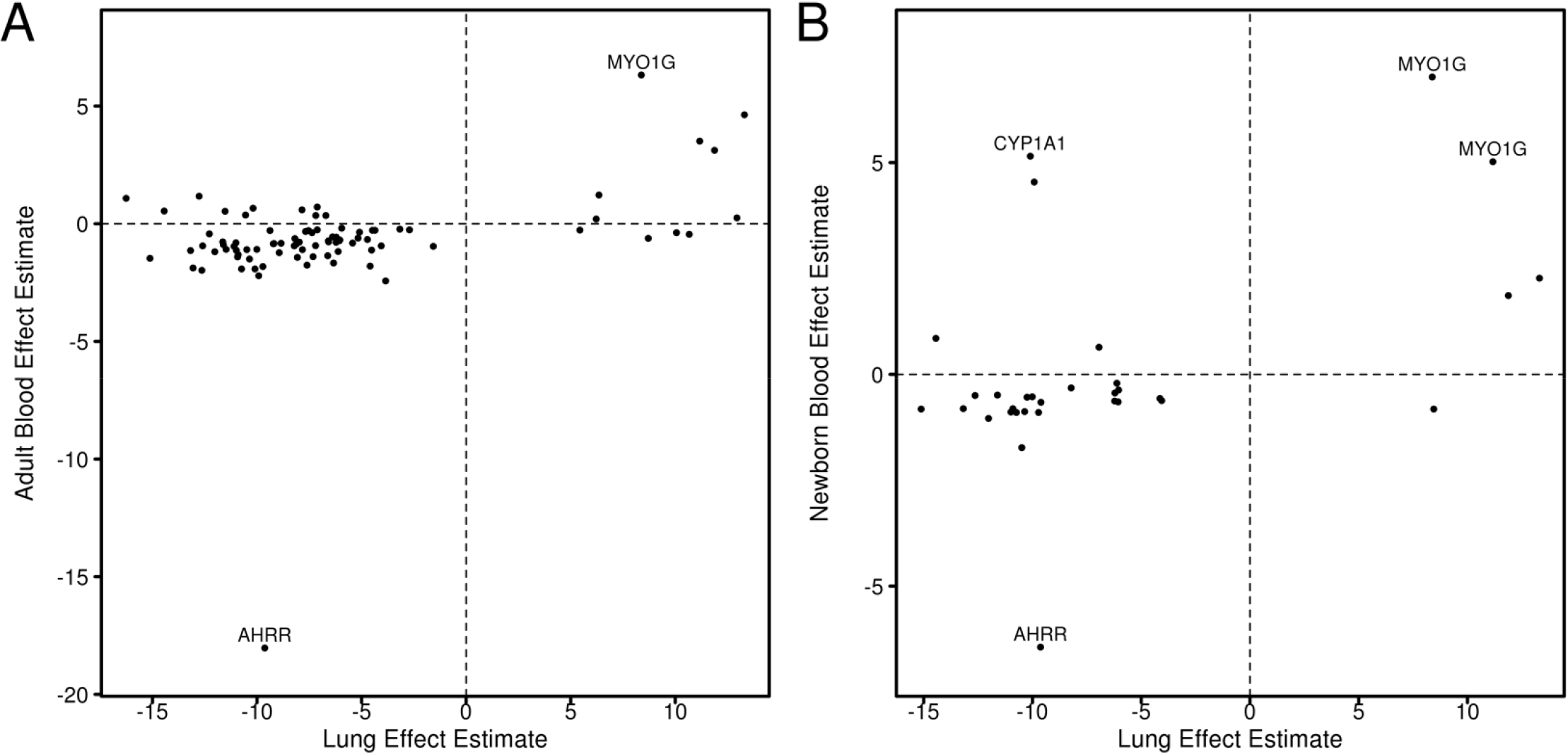
A. Effect estimates of CpG sites that are both associated with current smoking in adult blood (FDR<0.05) and current smoking status in lung adenocarcinoma (P<10^−4^) relative to never smokers. B. Effect estimates of CpG sites that are both associated with maternal smoking in newborn blood (FDR<0.05) and current smoking status in lung adenocarcinoma (P<10^−4^) relative to never smokers

CpG sites differentially methylated in relation to former smoking in lung adenocarcinoma were enriched for significant adult blood signals at a lower level than in lung of current smokers. CpG sites meeting the *P* threshold of 10^−4^ in lung adenocarcinoma from recent former smokers (≤15 years, n=193 sites) were enriched for the adult blood smoking signature (*P* = 1.4×10^−18^), with 41 sites overlapping with FDR significant CpGs in adult blood. Effect estimates between lung and adult blood for those 41 sites had Pearson correlation of 0.44 (**Supplementary Figure 6**). Top CpGs in lung adenocarcinoma from former smokers (>15 years, n=119 sites) were also enriched for significant signals in adult blood, with 12 overlapping sites (*P*=2.8×10^−3^). These former smoker lung adenocarcinoma and adult blood effect estimates for those 12 sites had correlation of 0.75.

We checked whether the sites associated with smoking in both adult blood and lung adenocarcinoma were consistent across exposure groups (current, recent former, longer former) (**Supplementary Figure 7**). Of the 86 sites overlapping between current smoker lung and adult blood, eight also had *P*<10^−4^ in the lung of both categories of former smokers.

#### Lung adenocarcinoma smoking DNA methylation signature is enriched for newborn blood smoking DNA methylation signature

We similarly compared the individual CpG site associations between current smoking in adenocarcinoma and maternal smoking in newborn blood. There were 5,936 CpG sites FDR associated with maternal smoking in newborns that were represented in the lung adenocarcinoma dataset. The differentially methylated CpGs in lung were enriched for signals in newborn blood, across multiple thresholds for lung significance (**Figure 1**). We again discuss results specific to the lung threshold of *P*<10^−4^. Among these CpG sites distinguishing current smokers versus never smokers in lung adenocarcinoma at *P*<10^−4^ (n=296 sites), we also observed enrichment for the newborn blood smoking exposure associations (*P* = 1.2×10^−19^). This corresponded to an observed overlap of 32 sites, relative to an expected overlap of 3.9 sites. Lung and newborn blood effect estimates for the 32 overlapping sites had Pearson correlation of 0.48 (**Figure 2B**) and 71.9% of those sites had reduced methylation with smoke exposure in both lung and newborn blood. As in the case of adult blood and lung adenocarcinoma, cg05575921 (*AHRR*) showed reduced methylation with smoking in newborns (6.4%). Two CpGs in *MYO1G* had some of the highest differences in methylation in both newborn blood and lung: cg04180046 (11.2% in lung, 5.0% in newborn blood) and cg12803068 (8.4% in lung, 7.0% in newborn blood). In *CYP1A1*, cg18092474 had one of the largest reduction methylation in relation to smoking in lung adenocarcinoma (10.1%), and was one of the sites with largest increased methylation in newborn blood (5.2%). Sites overlapping between lung adenocarcinoma and newborn blood were distributed among genomic regions in relation to CpG islands that differed from the full array distribution (G-test of goodness of fit *P*=1.7×10^−9^). We observed 71.9% of overlapping lung and newborn blood sites were in shore regions compared to 23.3% in the full array (*P*=3.5×10^−10^), 3.1% in open sea compared to 35.7% (*P*=2.6×10^−4^), 18.8% in islands compared to 31.6% (*P*=0.17), and 6.3% in shelf regions compared to 9.4% (*P*=0.74). Out of the 32 sites associated with sustained smoking in newborn blood at FDR < 0.05 and with current smoking in lung adenocarcinoma at *P*<10^−4^, 27 (84.4%) were similarly FDR significant in adult blood.

We tested for overlap between the lung adenocarcinoma former smoker associations with the newborn blood sites. Top CpGs (*P*<10^−4^, n=193 sites) in lung adenocarcinoma of former smokers (≤15 years) were enriched for smoking signals in newborn blood, with 28 sites overlapping with the significant newborn blood CpGs (*P* = 7.7×10^−21^). Effect estimates of the 28 sites had a correlation of 0.25. The newborn blood smoking signature had 9 CpGs in common with top CpGs (*P*<10^−4^, n=119 sites) in lung adenocarcinoma from former smokers (>15 years) (*P* = 2.8×10^−5^). These effect estimates had a correlation of 0.42. Of the 32 sites overlapping between current smoker lung and newborn blood, six also had *P*<10^−4^ in each of the two categories of former smokers in lung. These six sites were also FDR significant in adult blood. Among those common sites, four of the CpG sites (cg19089201, cg22132788, cg12803068 and cg04180046) are annotated to *MYO1G*, cg18092474 is annotated to *CYP1A1* and cg13985198 is annotated to *SNORD46, RPS8*, and *SNORD38A*.

### Sensitivity analyses

As a sensitivity analysis, enrichment tests were also performed for blood thresholds of *P*<10^−4^ as opposed to FDR<0.05. In adult blood 9,175 CpGs met this threshold, and in newborn there were 3,094 CpGs. Patterns of enrichment were similar to those seen using FDR significance criteria in blood (**Supplementary Figure 8**). Findings were also robust to varied *P* thresholds in lung adenocarcinoma CpGs. We allowed *P* cutoffs in lung associated CpGs to vary from 1.0 to 10^−10^. (**Figure 1**). When using FDR<0.05 cutoffs in lung results, 66 CpGs met the criteria in lung of current smokers, with 30 sites overlapping with FDR significant adult blood CpGs (P=1.2×10^−24^) and 14 sites overlapping with newborn blood CpGs (P=1.6×10^−13^). In lung of former smokers (≤15 years) 19 CpGs were FDR significant, with 14 overlapping with the adult blood signal (P=2.3×10^−16^) and 10 overlapping with newborn blood signal (P=1.1×10^−14^). In lung of former smokers (>15 years) 14 CpGs were FDR significant, with 4 overlapping with adult blood signal (P=1.8×10^−3^) and 3 overlapping with newborn blood signal (P=7.2×10^−4^).

### Gene ontology of sites uniquely associated with smoking in lung and not blood tissues

We investigated whether lung-specific DNA methylation changes in response to smoke exposure could provide insight to the biology of lung adenocarcinoma. Although the differentially methylated CpGs in lung adenocarcinoma were enriched for smoking signals in blood, we also identified several CpG sites in lung adenocarcinoma that were not implicated in either blood meta-analyses, both of which had a much larger sample size than the present study. There were 36 sites in current smoker lung that were FDR significant, 4 sites in former smokers (≤15 years), and 10 sites in former smokers (>15 years), that were not also FDR significant in adult blood or newborn blood (**Supplementary Table 3**).

To perform a pathway analysis on lung-specific sites, we used the relaxed threshold of *P*<10^−4^ to include a sufficient number of sites. The CpG sites that reached *P*<10^−4^ in lung adenocarcinoma of current smokers were annotated to 224 genes, and of those 65 were not implicated either the meta-analyses in blood. No biological pathways these 65 genes are involved in were enriched at an FDR significant level. The top pathways (P<0.01) include RNA and aromatic compound catabolic processes, and protein targeting (**Supplementary Table 4**). Many genes, especially ribosomal subunit genes, were involved in multiple pathways (**Figure 3**).

**Figure 3.**
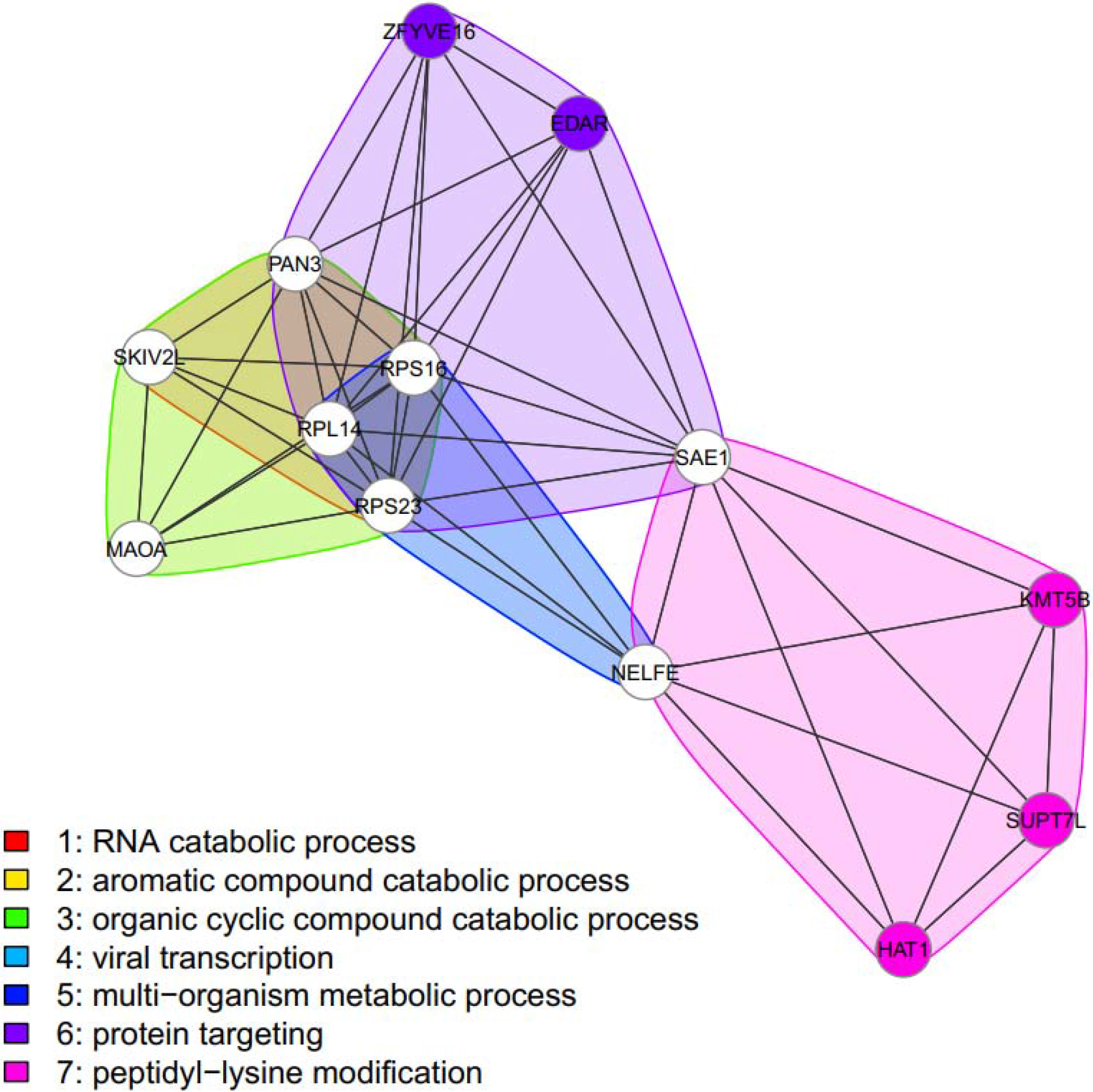
Network diagram of biological pathways enriched for genes with smoking-associated differentially methylated CpG sites meeting significance thresholds in lung tissue and not in blood meta analyses. Pathways with *P*<0.01 shown (none FDR significant). Nodes represent genes. Nodes are connected to other genes found in the same pathways. Color of nodes represent which pathway genes are found in, colored white if genes are in multiple pathways.

As a comparator, we also performed pathway analysis on overlapping lung adenocarcinoma-blood sites. A total of 78 genes were implicated by CpG sites that had *P*<10^−4^ in lung of current smokers and were also FDR significant in both adult and newborn blood. Several FDR significant gene ontologies for the common lung adenocarcinoma and blood smoking genes included many of the same or similar pathways, such as protein targeting to membrane and mRNA catabolic process. Other pathways enriched in the genes in common include translational initiation and various metabolic processes (**Supplementary Table 5**). Ribosomal subunit genes were also involved in most of the pathways. Genes found in both the top lung sites and blood sites include *AHRR*, *CYP1A1*, *CYP1B1*, *MYO1G*, several small nucleolar RNA genes, ribosomal subunit genes, and others (**Supplementary Table 6**).

### DISCUSSION

Tobacco smoking or exposure to tobacco smoke has been consistently associated with altered DNA methylation in blood measured across the life course (3, 5). Here, we identify DNA methylation alterations in lung adenocarcinoma associated with smoking and find both concordance, and discordance, between these smoking associated DNA methylation alterations and those previously reported in newborn and adult blood samples. These findings provide confirmatory evidence of a consistent smoking-associated DNA methylation signature across tissues. We also observed unique smoking DNA methylation association in lung adenocarcinoma tissue, suggesting there is a significant tissue specific component in epigenetic alterations in response to tobacco smoke. Interestingly, when stratifying analyses by recency of smoking cessation, the highest enrichment for smoking related DNA methylation changes previously observed in adult and newborn blood were seen in differentially methylated CpGs in lung adenocarcinoma of current smokers. These results support existing evidence that some DNA methylation alterations related to tobacco smoke exposure are attenuated with quitting time, and that the effects of tobacco smoke exposure in lung cancer tissue are not permanently mitotically heritable. These results, however, should be interpreted in light of the high rates of epigenetic dysregulation in tumors (20), where cells are rapidly cycling and there is persistent dysregulation of the epigenetic machinery. Epigenetic alterations in lung cancers could be caused directly by tobacco smoke exposure or reflect epigenetic changes as a result of cancer progression. There is a possibility that non-diseased, rather than cancerous, lung tissue from smokers would more closely reflect the epigenetic signature associated with tobacco smoke exposure in blood. For example, a recent study of epigenetic alterations identified seven significantly differentially methylated CpG sites in non-tumor lung tissue from smokers, five of which were also previously found to be differentially methylated in smoker’s blood (7).

We, and others, have identified smoking associated DNA methylation alterations in genes which have been identified to have functional roles relative to lung cancer initiation and progression. Epigenetic alterations may drive the formation of lung cancer by sensitizing cells to KRAS mutation (21). *AHRR* was hypomethylated with smoking in both lung cancer and blood, and hypomethylation of *AHRR* is associated with future lung cancer after adjustment for smoking (22), as well as low lung function, decline in lung function, and respiratory symptoms (23). While this may be explained, in part, by measurement error in self-reported smoking combined with *AHRR* methylation being an excellent quantitative biomarker of lifetime smoking behavior that captures this exposure better than questionnaires (24), smoking related reduced methylation in *AHRR* could play a role in pathogenesis. While a role for *MYO1G* in lung carcinogenesis has not been established, we also identified concordant methylation of CpG sites in *MYO1G* associated with smoking in lung cancer and both newborn and adult blood. Interestingly, siRNA knockdown of *MYO1G* in multiple cancer cell lines increased cell death and decreased autophagic flux, a process dysregulated in many human disorders (25). Whether *MYO1G* methylation in the lung is simply a biomarker of smoke exposure or has a functional role in cancer development remains to be determined. Intriguingly, while *CYP1A1* is hypermethylated relative to smoke exposure in newborn blood, we found a CpG site upstream of the *CYP1A1* transcription start site to be hypomethylated in lung cancer adenocarcinoma relative to smoking. *CYP1A1* polymorphisms have been linked to lung cancer risk, particularly when in combination with tobacco smoke (26–28), pointing to this gene’s important role in tobacco smoke toxicant metabolism and lung cancer etiology. Interestingly, the promoter of *CYP1A1* in normal lung tissue has been found to be hypermethylated in smokers (29), similar to the findings reported newborn blood, but not in lung cancer tissue or adult blood. However, the *CYP1A1* annotated CpGs hypomethylated in lung adenocarcinoma were also observed to have decreased methylation in adult blood findings. Determining whether the methylation changes in lung cancer associated with smoking are drivers or passengers of the carcinogenic progression will be essential to understand the clinical impact of these alterations.

The present study had several weaknesses. Despite a large number of lung adenocarcinoma cancer cases analyzed here (n=390), the power is much lower than in the two larger meta-analysis of adult and newborn smoking. The smoking-associated DNA methylation changes in the lung were not as numerous as those in adult and newborn blood which is likely to be consequence of the weaker power. Second, our lung tissue observations were derived from tumor tissues, which introduces the possibility that apparent smoking associated DNA methylation changes were a consequence of disease. Tumorigenesis is frequently associated with *de-novo* DNA methylation changes with enormous selection pressures on cells (30). Our observations demonstrate either smoking associated changes are able to survive tumorigenesis in lung adenocarcinoma, or they are recapitulated in the disease process, both suggesting they are of high biological importance in this tissue. The current study may also be repeated in non-diseased lung tissue as paired smoking status and DNA methylation measures become available. Further, tumors are heterogeneous mixtures of cell types, potentially also containing blood cells, each of which will have its own epigenetic profile. Recent work has shown that even within blood, some of the methylation changes associated with smoking are cell type specific (31). This, and future, studies would also be strengthened by the concurrent measurement of DNA methylation profiles of blood and a target tissue in the same individual. Determining whether smoking related epigenetic alterations in lung cancer are persistent across all cells or are relegated to individual cell types, with functional regulatory roles, will be an exciting future direction of research.

Integrated analyses of exposure related epigenomic signatures across tissue types represents a powerful approach to disentangle systemic and tissue specific alterations due to exogenous exposures. Here, for example, we identified a highly significant overlap in altered CpGs in lung tumor and blood due to smoking, providing support for the hypothesis that tissue types typically assayed by epidemiologists, such as blood, can provide relevant information about epigenetic alterations in target tissues, such as lung. The NIEHS Toxicant Exposures and Responses by Genomic and Epigenomic Regulators of Transcription (TaRGET) II consortium is currently formally testing this hypothesis across a range of exposures, tissue types, and epigenomic marks in mice (32). We also, however, identified many CpG sites altered related to smoking in lung cancer tissue, which were not previously reported in studies in blood (**Supplementary Table 3**). These CpGs not previously reported in blood likely represent unique effects in lung. If these CpGs were also differentially methylated in blood, the higher powered meta-analyses would likely have discovered them as well. Pathway analyses identified that these lung cancer specific alterations were enriched in genes and pathways involved in RNA catabolism, protein targeting, and transcription. Many ribosomal subunit proteins were represented in the top lung adenocarcinoma findings, many of which were also found in blood. The prevalence of these genes in pathway findings may be related to cancer, as ribosomal proteins often show increased expression in such cases (33). Understanding the tissue specific epigenomic signatures related to exposure may be able to identify etiologic agents in tumor development, critically impacting prevention and treatment.

## Acknowledgements

Dr. London is supported by the Intramural Research Program of the NIH, National Institute of Environmental Health Sciences (ZO1 ES49019). Mr. Dou and Dr. Bakulski were supported by grants from the National Institute of Environmental Health Sciences and the National Institute of Aging (R01 ES025531; R01 ES025574; and R01 AG055406). Dr. Colacino is supported by grants from the National Institute of Environmental Health Sciences (R01 ES028802 and P30 ES017885).

## REFERENCES

1. Anand P, Kunnumakkara AB, Sundaram C, Harikumar KB, Tharakan ST, Lai OS, et al. Cancer is a preventable disease that requires major lifestyle changes. Pharm Res. 2008 Sep;25(9):2097–116.

2. Cancer Genome Atlas Research N. Comprehensive molecular profiling of lung adenocarcinoma. Nature. 2014 Jul 31;511(7511):543–50.

3. Joubert BR, Felix JF, Yousefi P, Bakulski KM, Just AC, Breton C, et al. DNA Methylation in Newborns and Maternal Smoking in Pregnancy: Genome-wide Consortium Meta-analysis. Am J Hum Genet. 2016 Apr 07;98(4):680–96.

4. Reese SE, Zhao S, Wu MC, Joubert BR, Parr CL, Håberg SE, et al. DNA Methylation Score as a Biomarker in Newborns for Sustained Maternal Smoking during Pregnancy. Environ Health Perspect. 2017 Apr;125(4):760–6.

5. Joehanes R, Just AC, Marioni RE, Pilling LC, Reynolds LM, Mandaviya PR, et al. Epigenetic Signatures of Cigarette Smoking. Circ Cardiovasc Genet. 2016 Oct;9(5):436–47.

6. Shenker NS, Polidoro S, van Veldhoven K, Sacerdote C, Ricceri F, Birrell MA, et al. Epigenome-wide association study in the European Prospective Investigation into Cancer and Nutrition (EPIC-Turin) identifies novel genetic loci associated with smoking. Hum Mol Genet. 2013 Mar;22(5):843–51.

7. Stueve TR, Li WQ, Shi J, Marconett CN, Zhang T, Yang C, et al. Epigenome-wide analysis of DNA methylation in lung tissue shows concordance with blood studies and identifies tobacco smoke-inducible enhancers. Hum Mol Genet. 2017 Aug;26(15):3014–27.

8. Bibikova M, Barnes B, Tsan C, Ho V, Klotzle B, Le JM, et al. High density DNA methylation array with single CpG site resolution. Genomics. 2011 Oct;98(4):288–95.

9. Sandoval J, Heyn H, Moran S, Serra-Musach J, Pujana MA, Bibikova M, et al. Validation of a DNA methylation microarray for 450,000 CpG sites in the human genome. Epigenetics. 2011 Jun;6(6):692–702.

10. Chen YA, Lemire M, Choufani S, Butcher DT, Grafodatskaya D, Zanke BW, et al. Discovery of crossreactive probes and polymorphic CpGs in the Illumina Infinium HumanMethylation450 microarray. Epigenetics. 2013 Feb;8(2):203–9.

11. Triche TJ, Jr., Weisenberger DJ, Van Den Berg D, Laird PW, Siegmund KD. Low-level processing of Illumina Infinium DNA Methylation BeadArrays. Nucleic Acids Res. 2013 Apr;41(7):e90.

12. Ritchie ME, Phipson B, Wu D, Hu Y, Law CW, Shi W, et al. limma powers differential expression analyses for RNA-sequencing and microarray studies. Nucleic Acids Res. 2015 Apr 20;43(7):e47.

13. Smyth GK. Linear models and empirical bayes methods for assessing differential expression in microarray experiments. Stat Appl Genet Mol Biol. 2004;3:Article3.

14. Barfield RT, Almli LM, Kilaru V, Smith AK, Mercer KB, Duncan R, et al. Accounting for population stratification in DNA methylation studies. Genet Epidemiol. 2014 Apr;38(3):231–41.

15. Leek JT, Storey JD. Capturing heterogeneity in gene expression studies by surrogate variable analysis. PLoS Genet. 2007 Sep;3(9):1724–35.

16. Devlin B, Roeder K. Genomic control for association studies. Biometrics. 1999 Dec;55(4):997–1004.

17. Andrews SV, Ladd-Acosta C, Feinberg AP, Hansen KD, Fallin MD. “Gap hunting” to characterize clustered probe signals in Illumina methylation array data. Epigenetics Chromatin. 2016;9:56.

18. Phipson B, Maksimovic J, Oshlack A. missMethyl: an R package for analyzing data from Illumina’s HumanMethylation450 platform. Bioinformatics. 2016 Jan;32(2):286–8.

19. Supek F, Bošnjak M, Škunca N, Šmuc T. REVIGO summarizes and visualizes long lists of gene ontology terms. PLoS One. 2011;6(7):e21800.

20. Virani S, Colacino JA, Kim JH, Rozek LS. Cancer epigenetics: a brief review. ILAR J. 2012;53(3-4):359–69.

21. Vaz M, Hwang SY, Kagiampakis I, Phallen J, Patil A, O’Hagan HM, et al. Chronic Cigarette Smoke-Induced Epigenomic Changes Precede Sensitization of Bronchial Epithelial Cells to Single-Step Transformation by KRAS Mutations. Cancer Cell. 2017 Sep;32(3):360–76.e6.

22. Fasanelli F, Baglietto L, Ponzi E, Guida F, Campanella G, Johansson M, et al. Hypomethylation of smoking-related genes is associated with future lung cancer in four prospective cohorts. Nat Commun. 2015 Dec;6:10192.

23. Kodal JB, Kobylecki CJ, Vedel-Krogh S, Nordestgaard BG, Bojesen SE. AHRR hypomethylation, lung function, lung function decline and respiratory symptoms. Eur Respir J. 2018 Mar;51(3).

24. Valeri L, Reese SL, Zhao S, Page CM, Nystad W, Coull BA, et al. Misclassified exposure in epigenetic mediation analyses. Does DNA methylation mediate effects of smoking on birthweight? Epigenomics. 2017 03;9(3):253–65.

25. Groth-Pedersen L, Aits S, Corcelle-Termeau E, Petersen NHT, Nylandsted J, Jäättelä M. Identification of Cytoskeleton-Associated Proteins Essential for Lysosomal Stability and Survival of Human Cancer Cells. PLoS ONE. 2012 10/11 05/03/received 08/17/accepted;7(10):e45381.

26. Ji Y-N, Wang Q, Suo L-j. CYP1A1 Ile462Val Polymorphism Contributes to Lung Cancer Susceptibility among Lung Squamous Carcinoma and Smokers: A Meta-Analysis. PLOS ONE. 2012;7(8):e43397.

27. Shaffi SM, Shah MA, Bhat IA, Koul P, Ahmad SN, Siddiqi MA. CYP1A1 polymorphisms and risk of lung cancer in the ethnic Kashmiri population. Asian Pacific journal of cancer prevention: APJCP. 2009 Oct-Dec;10(4):651–6.

28. Song N, Tan W, Xing D, Lin D. CYP 1A1 polymorphism and risk of lung cancer in relation to tobacco smoking: a case-control study in China. Carcinogenesis. 2001;22(1):11–6.

29. Anttila S, Hakkola J, Tuominen P, Elovaara E, Husgafvel-Pursiainen K, Karjalainen A, et al. Methylation of Cytochrome P4501A1 Promoter in the Lung Is Associated with Tobacco Smoking. Cancer Research. 2003;63(24):8623–8.

30. Tabassum DP, Polyak K. Tumorigenesis: it takes a village. Nat Rev Cancer. 2015 Aug;15(8):473–83.

31. Su D, Wang X, Campbell MR, Porter DK, Pittman GS, Bennett BD, et al. Distinct Epigenetic Effects of Tobacco Smoking in Whole Blood and among Leukocyte Subtypes. PLoS One. 2016;11(12):e0166486.

32. Wang T, Pehrsson E, Purushotham D, Li D, Zhang B, Lawson H, et al. The NIEHS TaRGET II Consortium and Environmental Epigenomics. Nature Biotechnology. 2018 2018.

33. Xie X, Guo P, Yu H, Wang Y, Chen G. Ribosomal proteins: insight into molecular roles and functions in hepatocellular carcinoma. Oncogene. 2018 Jan;37(3):277–85.

